# Effect of muscarinic blockade on the speed of attention shifting and learning

**DOI:** 10.1101/2024.05.08.593141

**Authors:** Alexander Thiele, Agnes McDonald Milner, Corwyn Hall, Lucy Mayhew, Anthony Carter, Sidharth Sanjeev

## Abstract

The study aimed to investigate to what extent blockade of muscarinic receptors affects the speed of endogenous versus exogenous attentional shift times, and how it affects learning induced improvements of attention shift times. Subjects viewed an array of 10 moving clocks and reported the time a clock indicated when cued. Target clocks were indicated by peripheral or central cues, including conditions of pre-cuing. This allowed assessing shift times when attention was pre-allocated, when peripheral cues triggered exogenous attention shifts, and when central cues triggered endogenous attention shifts. In study 1, each subject participated in 2 sessions (scopolamine/placebo), whereby the order of drug intake was counterbalanced across subjects, and subjects were blinded to conditions. Scopolamine/placebo was administered before a psychophysical experiment was conducted. In study 2, the effect of muscarinic blockade on learning induced improvements of attention shift times was investigated. Here scopolamine/placebo was administered immediately after the first (of two) psychophysical sessions, whereby a given subject either received scopolamine or placebo pills. Confirming previous results, we show that pre-cuing resulted in the fastest shift times, followed by exogenous cuing, with endogenous attentional shifts being slowest. Scopolamine application increased attentional shift times across all 3 conditions compared to placebo, but in a dose dependent manner. Additionally, blockade of muscarinic receptors immediately after the first session reduced learning dependent improvement of attention shift times. These results demonstrate that muscarinic receptors play an important role in attention shifting, and they contribute to learning of attention shifting.

## Introduction

The speed with which covert attention can be shifted between different locations, has been investigated using many different paradigms, dating back to experiments performed by Wilhelm Wundt using a complication clock apparatus (Wundt 1883). Orienting paradigms (Posner 1980; Yantis and Jonides 1990; Carlson et al. 2006), paradigms investigating attentional gating (Reeves and Sperling 1986) and visual search paradigms (Wolfe et al. 2000) have been used in different forms. These studies have demonstrated that two types of attention can be dissociated, namely bottom-up and top-down attention (Posner 1980; Duncan 1984; Posner and Petersen 1990; Theeuwes 1991; Carlson *et al*. 2006). Bottom-up attention is reflexive and triggered by salient unexpected stimuli in the external world (often peripheral cues in experimental situations), conversely top-down attention is wilful, and can be triggered by central cues (which often need to be interpreted in experimental situations). These two types of attention also differ in terms of how quickly they can be allocated (shifted) to stimuli of interest. Bottom-up attention shifts are faster than top-down attention shifts by ∼50-100ms (Posner 1980; Muller and Rabbitt 1989; Carlson *et al*. 2006; Chakravarthi and VanRullen 2011).

To what extent these types of attention depend on different neurotransmitter/neuromodulator systems is unknown (Thiele and Bellgrove 2018). Attentional selection is assumed to be driven by frontal and parietal cortex (Corbetta and Shulman 2002; Moore and Armstrong 2003). Parietal and frontal regions influence sensory processing directly via feedback, but also indirectly, via connections to cholinergic neurons in the basal forebrain that have ascending projections to sensory areas (Russchen et al. 1985; Sarter et al. 2005). Critically, ample evidence for the involvement of acetylcholine (ACh) in attentional modulation exists (Nelson et al. 2005; Robbins 2005; Sarter *et al*. 2005; Parikh et al. 2007; Furey et al. 2008). Disrupting cholinergic fibres originating in the basal forebrain, thereby depleting cortical areas of ACh, results in attentional impairments (McGaughy et al. 2002; Sarter *et al*. 2005). Attentional deficits in Alzheimer’s disease partly arise from cholinergic dysfunction (Nobili and Sannita 1997). Infusion of the muscarinic blocker scopolamine into macaque parietal cortex results in a dose-dependent increase in reaction times and a decrease in accuracy, when locations were exogenously cued, with the biggest effect for validly cued locations (Davidson et al. 1999; Davidson and Marrocco 2000). Muscarinic blockade also reduced performance in sustained visual attention task, which can be conceptualized as an endogenous attention task (Ellis et al. 2006), and reduces attentional modulation of cell responses in primary visual cortex and frontal eye field neurons in macaque monkeys under endogenously cued covert attention conditions (Herrero et al. 2008; Dasilva et al. 2019). These studies suggest that endogenous and exogenous attention are both dependent on muscarinic signalling, but a direct test has, to the best of our knowledge, never been done.

To test the role of muscarinic signalling in exogenously, endogenously, and pre-cued attention shifts, we used a variant of Wundt’s complication-clock apparatus (Wundt in Carlson *et al*. 2006). It involves participants watching a fixation point surrounded by moving clocks. After a clock is cued, participants shift their attention to that clock and record the position of the hand they perceived on the clock when the clock was cued. In study 1 participants performed the task twice on different days, either after having taken oral doses of scopolamine, or placebo pills.

Muscarinic receptors also play an important role in neuronal plasticity (McKenna et al. 1989; Metherate and Weinberger 1990; Weinberger and Bakin 1998; Barros et al. 2002) and learning (McGurk et al. 1988, 1991; Carli et al. 1997; Izquierdo et al. 1998; Thiel et al. 2002; Hasselmo 2006; Barker and Warburton 2009; Thiele 2013). A recent study has shown that attention shift times are affected by learning (*Thiele et al*., *Biorxiv*). We therefore also investigated the role of muscarinic signalling in the learning dependent improvement of attention shift times. Here an oral dose of either placebo or scopolamine was given to participants after the first experimental session, and they then performed a second experimental session on the following day.

We found that muscarinic blockade affected attention shift times in a dose dependent manner. No differences of the drug were found between endogenously, exogenously, or pre-cued attention shifts. Moreover, application of the muscarinic blocker scopolamine during the immediate consolidation period resulted in reduced improvements of attention shift times.

## Methods

### Participants

Study 1: Eighty-one participants took part in the experiment, where the effect of muscarinic blockade on attention shift times was investigated. This sample consisted of 34 males and 47 females, with an age range of 18-29 years, median 22.

Study 2: Twenty subjects took part in the experiment where the effect of muscarinic blockade on learning induced improvements of attention shift times was tested. The sample consisted of 20 students (10 male/10 female) from Newcastle University. Participants were selected based on opportunity sampling and had an age range of 19 – 22 years old, median 21.

All participants had normal or corrected-to-normal vision. They were required to sign a consent form to participate in the experiment. The consent form gave information about the task, the detailed procedures, the purpose of the study, how data would be used and anonymized. Here they also confirmed that they had not eaten for 2 hours before the experiment or drunk alcohol for 12 hours before the experiment, were not hungover at the day of the experiment and had abstained from using recreational drugs for >48h. Subjects were only included if they were at least 18 years old, not allergic to Kwell pills (active ingredient hyoscine hydrobromide, also known and referred to as scopolamine) or if they were taking any medication which should not be taken alongside Kwell pills. High blood pressure and pregnancy were also exclusion criteria. They were instructed not to drive or operate any motorised equipment for up to 6 hours after a session. Ethics were approved by the Faculty of Medical Sciences ethics committee at Newcastle University (01016_1/2015 to 01016_6/2023). Subjects had to perform the task twice, whereby each session had to be performed on separate days (at about the same time of a day, i.e. if a subject did session one in the morning, they performed session 2 on a different day, but also in the morning).

### Design

Two separate designs were used for the Study 1 and Study 2.

Study1: Assessing the effects of scopolamine on attention shift times. The design of the experiment was within-subject repeated measurements. Each participant carried out the experimental and placebo conditions in separate sessions. The two sessions were performed on different days, but at roughly the same time of day to control for effects of circadian rhythm on attention. They were informed prior to the experiments that in one of the two sessions they would receive either a dose of Kwell pills, or one/two placebo pills. Subjects were given the respective pills 45 mins before the start of the experiments, after which they were free to spend the next 45 minutes in whichever way they chose. Forty-one subjects were given a dosage of 300mg scopolamine, when the ‘drug’ session was selected, and forty subjects were given a dosage of 600mg scopolamine, when the ‘drug’ session was selected. Placebos were homeopathic arnica 6C. Forty of participants performed the experimental condition first and the remaining forty-one performed the placebo condition first, to counterbalance potential order (learning) effects.

Study 2: Assessing the effects of scopolamine on learning induced improvement of attention shift times: The design of the experiment was between-subject repeated measurements. Each participant carried out two separate sessions, whereby one group of subjects received two Kwell pills immediately after the first session, and the other group received the placebo pills immediately after the first session. A single dosage of 600mg was used in the subjects that received the drug treatment. Placebo treatment consisted of two unflavoured chewable glucose pills. Subjects were blinded to the condition. They were then free to leave the premises, to return at around the same time on the following day, when they performed the second session.

### Procedure

Before the experiment, participants were given the experiment information sheet and questionnaire via email. They had to fill out the pre-experiment questionnaire and email it to the experimenter. Upon arrival for the first session, they were given a consent form to read and sign. They were briefed again about the experiment, i.e. the psychophysical task in conjunction with muscarinic blockade. Any questions about the task were then answered. This was followed by a 30-trial practice session, where they familiarized themselves with task and obtained initial training.

The psychophysical task was as described in (Carlson *et al*. 2006). It consisted of 3 different attention conditions (pre-cue, exogenous and endogenous cuing), where subjects had to read the time on one of 10 possible clocks, when the relevant clock was cued (see below). Each condition occurred 50 times, randomly selected without replacement. Each trial started with a central fixation spot, which subjects were asked to fixate. Before the start of the trial the computer randomly assigned which clock would be cued on that given trial. In the pre-cue conditions the fixation spot appeared along with a central cue, a short line (length: 2 deg of visual angle (dva)), emanating from close to the fixation cross, pointing to the future relevant clock location. The cue was presented for 5 frames (50ms) after which it disappeared while the fixation spot stayed on. After 1 second, 10 clock faces (∼2.5 dva diameter) appeared simultaneously at an eccentricity of 7 dva on the screen. Each clock showed a single clock hand with a randomly allocated starting position (see figure 1 for an example). The clock hands rotated at 1Hz, i.e. one revolution per second. At a randomly selected time after the start of the trial (drawn from a uniform distribution from 100-1000ms after clock onset, steps of 10ms) one of the clocks would be cued (details below). Shortly after the cue (1000ms), the clocks were replaced by one central clock. Participants were required to indicate the position of the hand on the clock when it was cued by moving the hand of the second clock to the same position using the left/right arrow buttons of the keyboard. There were 3 different conditions in the task; (1) in the exogenous cue condition, the rim of the cued clock changed from black to red at the time of cuing for 5 frames (50ms). (2) In the endogenous cue condition, a line appeared emanating from the fixation point pointing towards one of the clocks (for 5 frames, i.e. 50 ms). (3) In the baseline (pre-cue) condition, a line appeared before the clock faces (as described above), indicating the position of the clock that would be cued. That specific clock was then cued during the trial as in the exogenous cuing condition. Figures 1 illustrates the progression of events in each of the 3 conditions (see also the videos in Carlson *et al*. 2006).

**Figure 1.**
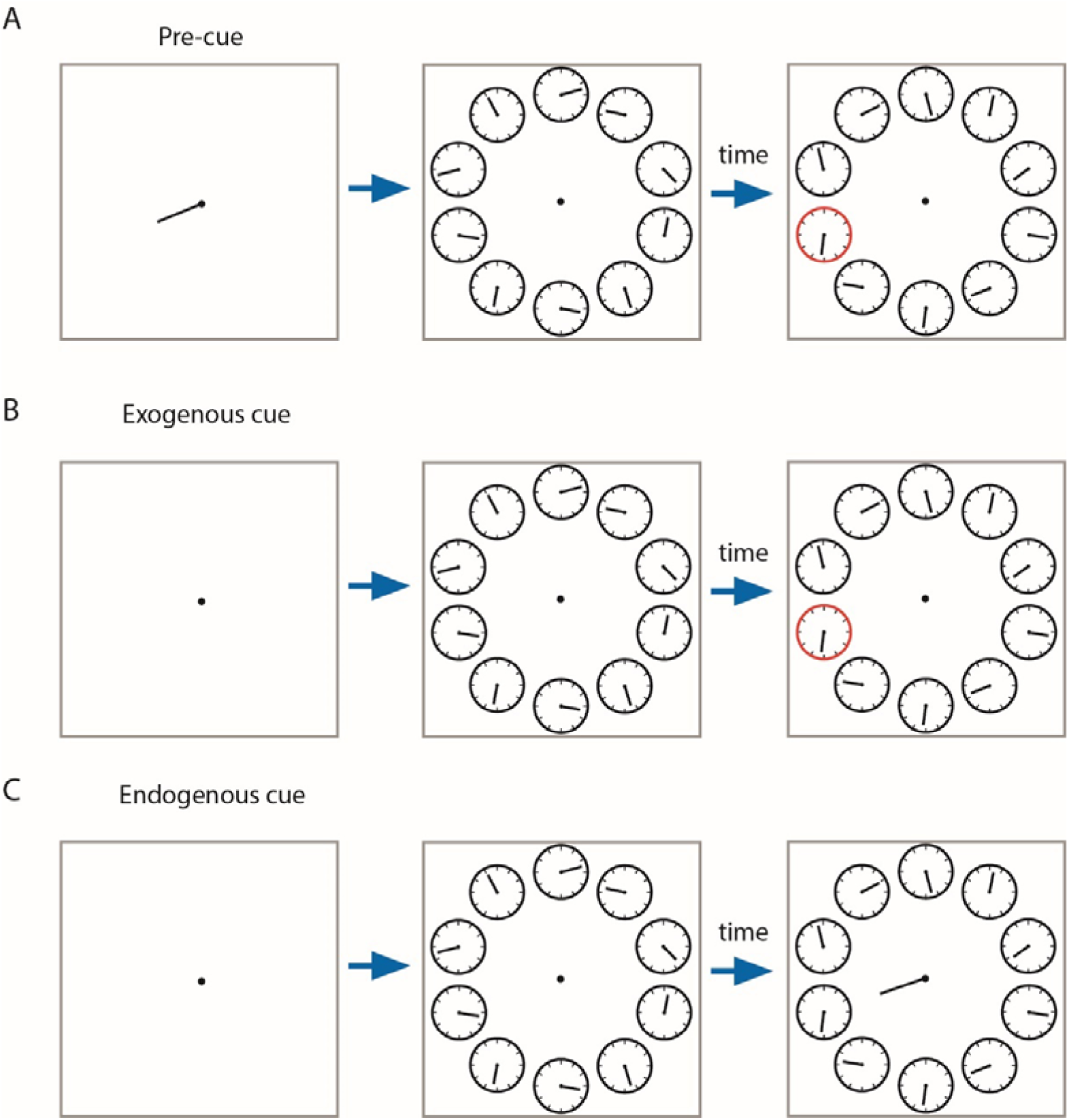
Experimental setup for the 3 attention conditions. A) The top row indicates the events that occur in the pre-cue condition, B) middle row the events in the exogenous condition, and C) bottom row the events that occur in the endogenous cuing condition. The blue arrow indicates time passing. This was fixed for the first period, but variable for the second period.

### Apparatus/ Materials

The experiment was run on windows PC running MATLAB 2015a (62-bit) (The MathWorks, Inc.) using PsychToolbox extentions (Brainard 1997). The stimuli were displayed on an Iiyama 22” monitor (100 Hz, 2,048 x 1536 resolution) that was controlled by the PC. Observers sat with their chin on a chin rest 60cm away from the monitor. Responses were collected from the keyboard, using the left and right arrow keys to move the clock hand and the space bar to confirm the participant’s observed time.

### Data Analysis

The latency between the real time of cueing and the time of cueing reported/perceived by the participant (i.e. the time indicated by the adjusted the clock hand) was calculated for each trial. This could result in latencies of 0-999ms (one full revolution of the clock hand) in principle. However, latencies that exceeded 900 ms were calculated as latency-1000. This cutoff latency was accepted as it allows for some random perceptual/memory errors to be distributed around the actual time that was present at cuing. It resulted in all final latencies to range from -99ms to 900ms. The mean latency for the 3 cuing and the 2 experimental (drug/no drug) conditions were then calculated for each participant.

To investigate whether scopolamine affected attention shift times a mixed model ANOVA was used. This was based on subject means calculated for the three cuing conditions (3 cuing conditions, within subject repetition), experimental condition (drug/no drug, within subject repetition), drug dosage (300mg/600 mg, between subject factor), or the order of the experimental condition (scopolamine or placebo in first session, between subject factor).

To investigate whether scopolamine affected the learning of attention shift times a mixed model ANOVA was also used, but with different factors. This was based on subject means calculated for the three cuing conditions (3 cuing conditions, within subject repetition), and the experimental condition (drug/no drug, between subject factor).

## Results

We will first describe the results regarding the effects of muscarinic blockade on attention shift times, followed by describing the effects on learning induced improvements of attention shift times.

### Effects of muscarinic blockade on attention shift times

Eighty-one participants performed the study, whereby 41 participants took the placebo before the first session, and 40 participants took the Kwell pill(s) before the first session (order of experimental conditions). Forty-one subjects took a dose of 300mg in the drug condition (21 subjects took the placebo first, the remaining subjects took the drug first), and the remaining 40 subjects took a dose of 600mg in the drug condition (20 subjects took the placebo first, the remaining subjects took the drug first). Confirming previous results (Carlson *et al*. 2006), we found that shift times were fastest in the pre-cue condition, followed by the exogenous cuing, and slowest in the endogenous cuing condition (Figure 2).

**Figure 2.**
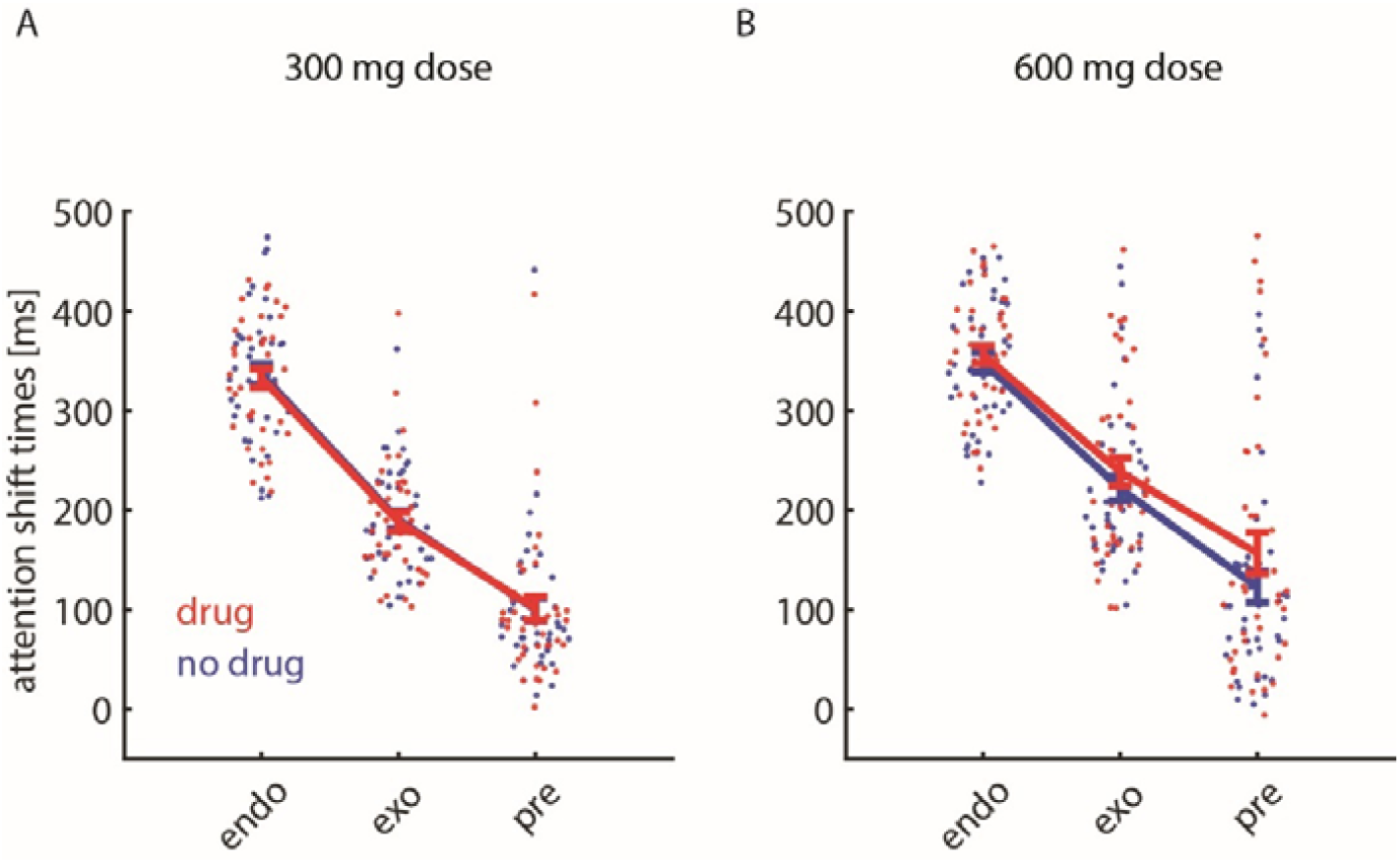
Attentional shift times for the 3 attention conditions when subjects were not under the influence of scopolamine (blue) and when they were under the influence of scopolamine (red). Cuing conditions are indicated along the x-axis, mean attentional shift times along with S.E.M are indicated along the y-axis. A) data for subjects in the 300mg drug study (n=41), B) data for subjects in the 600mg drug study (n=40).

The mixed model ANOVA revealed that there was a significant main effect of cue condition, a significant main effect of drug condition, a significant effect of drug level (table 1). Additionally, there were significant interactions between order and drug (y/n), as well as between drug (y/n) and drug-level (300/600 mg) (table 1).

**Table 1.** Mixed model ANOVA details. *Factor* indicates the parameter of interest, *df* shows the degrees of freedom, F and p give the *F*- and *p*-values respectively. * symbol denotes interaction between factors. Significant factors (and interactions) have been highlighted in grey, for ease of reading.

| Factor | df | F | p |
| --- | --- | --- | --- |
| Intercept (subject effects) | (1,462) | 1130.8 | <0.001 |
| cue-type | (2,462) | 784.7 | <0.001 |
| drug (y/n) | (1,462) | 4.1 | 0.044 |
| order | (1,462) | 2.7 | 0.099 |
| drug-level (300/600mg) | (1,462) | 7.0 | 0.008 |
| order*drug (y/n) | (1,462) | 11.6 | <0.001 |
| order*cue-type | (2,462) | 0.9 | 0.399 |
| drug (y/n)* cue-type | (2,462) | 1.0 | 0.363 |
| order*drug-level (300/600mg) | (1,462) | 0.1 | 0.819 |
| drug (y/n)*drug-level (300/600mg) | (1,462) | 6.5 | 0.011 |
| cue-type*drug-level (300/600mg) | (2,462) | 2.3 | 0.102 |
| order*drug(y/n)*cue-type | (2,462) | 0.0 | 0.954 |
| order*drug(y/n)*drug-level (300/600mg) | (1,462) | 0.1 | 0.793 |
| order*cue-type*drug-level (300/600mg) | (2,462) | 1.7 | 0.192 |
| drug(y/n)*cue-type*drug-level (300/600mg) | (2,462) | 0.7 | 0.514 |
| order*drug(y/n)*cue-type*drug-level (300/600mg) | (2,462) | 0.6 | 0.553 |

Scopolamine intake increased attentional shift times for all 3 cue types, with no interaction effect between drug (y/n) and cue types (Table 1, and Figure 2). This indicates that the negative effect of muscarinic blockade (the slowing) on attentional shift times does not depend on the cuing condition (pre-, exogenous or endogenous). Drug level significantly affected attentional shift times, whereby visual inspection (figure 2) suggests that 300mg hardly resulted in a slowing, while 600mg caused larger slowing. This impression is corroborated by analysing drug-levels separately. The 300mg dose did not affect attentional shift times (main effect of drug: p=0.689, F(1,234)=0.2, mixed model ANOVA), while the 600 mg dose significantly affected attentional shift times (main effect of drug: p=0.004, F(1,228)=8.7, mixed model ANOVA). Based on figure 2 it appears that slowing is most profound for the pre-cue condition, when subjects were exposed to a 600 mg dose, but the lack of an interaction effect between cue-type and drug-level argues against that idea.

The order of placebo/drug sessions itself was not significant (p=0.099, mixed model ANOVA, table 2) which might indicate that shift times did not differ between the first and the second session. However, there was a significant interaction between drug (y/n) and order (p<0.001, table 1). Drug application generally increased attentional shift times, and hence any learning (improvement) between session, might be concealed in the group that was under the influence in the second session. This argument is supported by the significance of the drug* order interaction (mixed model ANOVA: p=0.008, table 2). Subjects usually had faster shift times in the second compared to the first experimental session, (Figure 3).

**Table 2.** Mixed model ANOVA details relating to study 2. *Factor* indicates the parameter of interest, *df* shows the degrees of freedom, *F* and *p* give the F- and p-values respectively. * symbol denotes interaction between factors. Significant factors (and interactions) have been highlighted in grey, for ease of reading.

| Factor | df | F | P |
| --- | --- | --- | --- |
| Intercept (subject effects) | (1,108) | 265.3 | <0.001 |
| cue-type | (2,108) | 182.1 | <0.001 |
| drug (y/n) | (1,108) | 0.8 | 0.368 |
| day | (1,108) | 3.2 | 0.077 |
| drug (y/n)* cue-type | (2,108) | 1.5 | 0.232 |
| day* cue-type | (1,108) | 0.1 | 0.896 |
| drug (y/n)*drug | (1,108) | 7.2 | 0.009 |
| day*drug (y/n)*cue-type | (2,462) | 0.4 | 0.663 |

**Figure 3.**
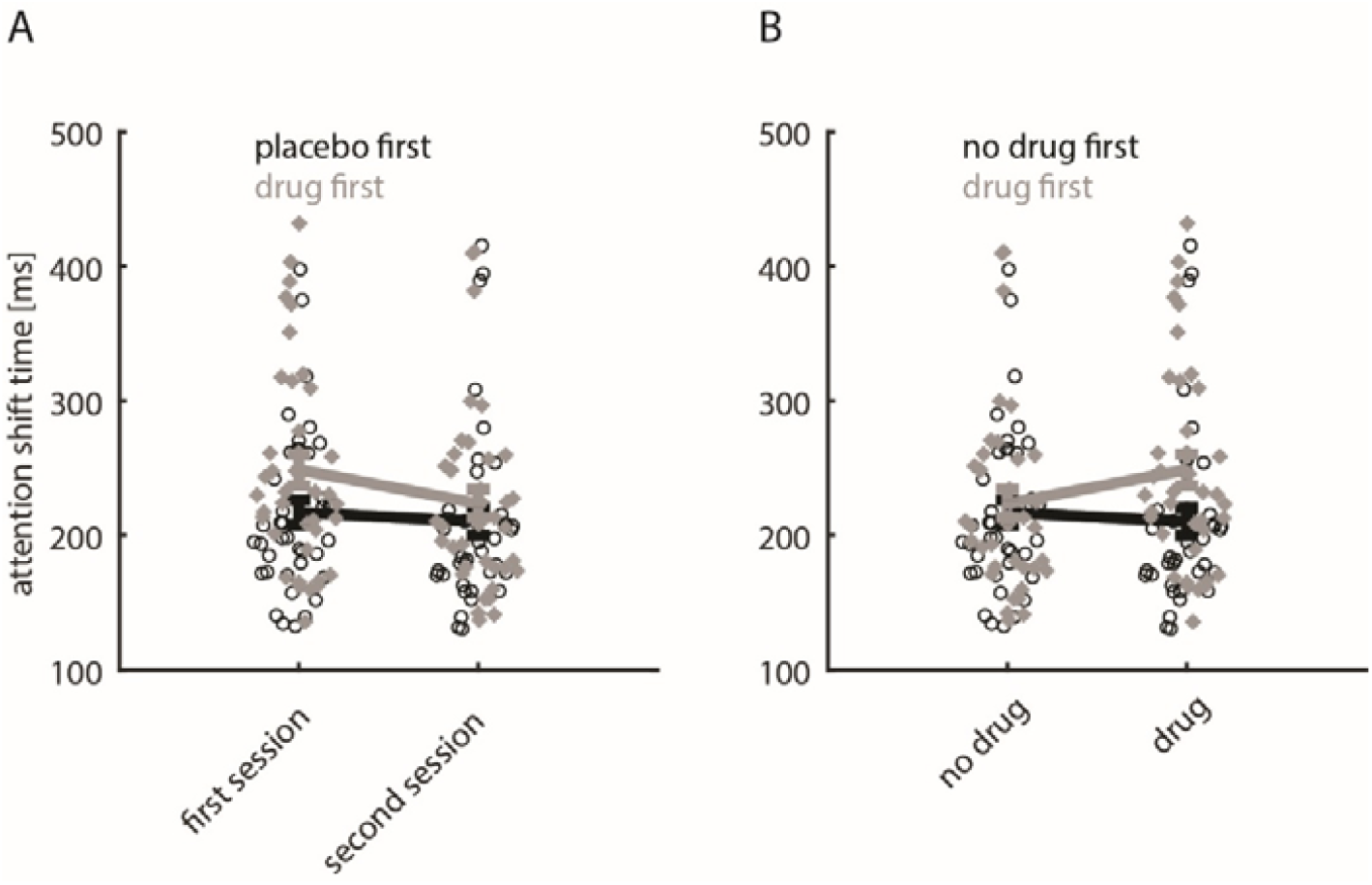
Average attentional shift times (across the 3 conditions). A) Shift times as a function of session (first, second) and as a function of whether they were under the influence of scopolamine (drug first, black) in the first session. B) Shift times as a function of whether they were under the influence of scopolamine or not, separated whether scopolamine was given in the first session (grey, drug first) or whether placebo was given in the first session. Note that for grey data points in B that are located on the right correspond to participants who took the drug in the first session, while grey data points on the left correspond to the same subjects who participated under placebo conditions (hence the no drug label) in the second session. Session type is indicated along the x-axis, mean attentional shift times along with S.E.M are indicated along the y-axis.

Figure 3A shows that subjects’ shift times were comparatively slow when they were under the influence of scopolamine in the first session, while shift times were faster in the second session when they were not under the influence of scopolamine. Separating these effects for different drug levels shows this effect more clearly (Figure 4).

**Figure 4.**
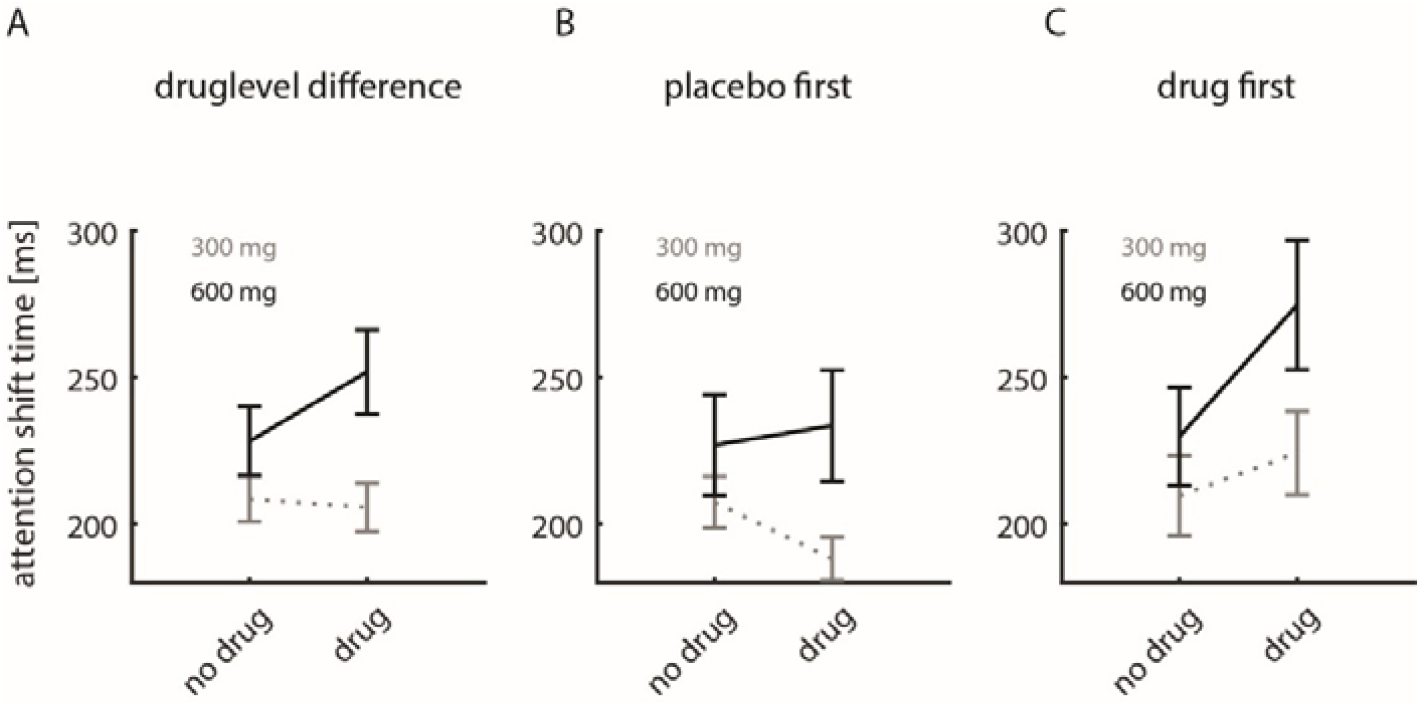
Average change in attentional shift times for different drug level (300mg vs. 600 mg). A) Difference in average attention shift times between drug and no drug conditions, independent of the order of drug/placebo session. B) Change in attention shift times for sessions where the placebo was taken in the first session. C) Change in attention shift times for sessions where the drug (scopolamine) was taken in the first session.

Figure 4A shows that shift times were similar between no drug (placebo) and drug session when a dose of 300mg was given in the drug session. When a 600mg dose was given in the drug sessions average attention shift times were increased compared to placebo session. This shows the drug-level effect previously described (see also figure 2 and table 1). When placebo was taken in the first session and 300mg scopolamine were taken in the second session, subjects had faster attention shift times in the second session (figure 4B). Conversely, when 600mg scopolamine were taken in the second session, attention shift times were slightly increased (figure 4B). This pattern might suggest that low dosages of scopolamine decrease attention shift times, while high dosages increase them. However, it becomes clear that this is not the case, when analysing the data from sessions where the drug was taken first (figure 4C). In this case drug sessions always resulted in slower attention shift times, when compared to placebo sessions. The data from figure 4B and C visually underpin the significant order* drug (y/n) interaction effect (table 1). An order effect would demonstrate that subjects show some form of learning between sessions. While there was no main effect of order present, the interaction effect between order and drug (y/n) demonstrates that the learning (order effect) depends on whether or not scopolamine was taken in the first session. The design involved in the study described so far does not allow to determine the contribution of muscarinic receptors to learning directly, as performance in one of the two session was always affected by muscarinic blockade, in addition to the possible effect on consolidation that takes place between session.

We therefore performed an additional study (Study 2, methods), where performance within a session was always under control conditions (neither drug nor placebo was taken). The independent variable was whether subjects received scopolamine (group 1, n=10, 2*300mg Kwell pills) immediately after the first session (to potentially affect consolidation), or whether they received placebo after the first session (group 2, n=10, 2 unflavoured glucose pills).

Figure 5A shows that the average attention shift times were reduced in subjects who took the placebo pills after the first session for all 3 cue-types. This was not the case for the group of subjects who took scopolamine after the first session. To determine which factors significantly affected attention shift times we performed a mixed-model ANOVA (table 2). Cue-type significantly affected performance, drug (y/n) had no effect on its own, there was a mild trend that day (session 1/2) had an effect on shift times, but critically there was a significant interaction between drug (y/n) and day (session 1/session 2).

**Figure 5.**
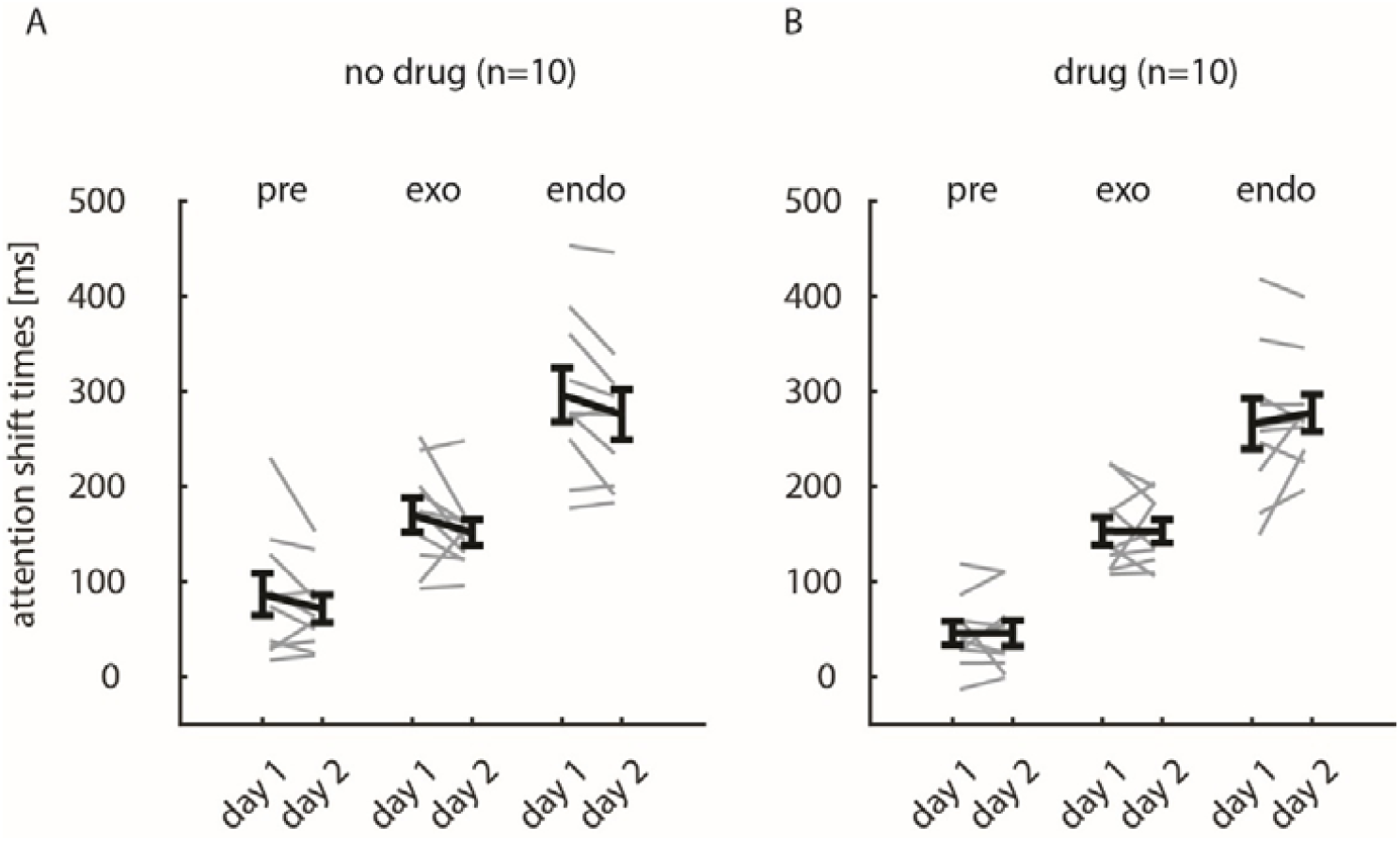
Average attentional shift times (across the 3 conditions) as a function of session (day 1, day 2). A) Data from subjects who took placebo after the first session. B) Data from subjects who took the drug (600mg scopolamine) after the first session. Mean attentional shift times along with S.E.M are indicated along the y-axis in black, grey lines show means for individual subjects.

To further investigate the interaction between drug (y/n) and day, we subtracted the attention shift times in session 2 from the attention shift times in session 1. This indicates whether a systematic change occurred between sessions, and whether this differed for the two treatment groups. The distribution of differences for the two groups is shown in figure 6.

**Figure 6.**
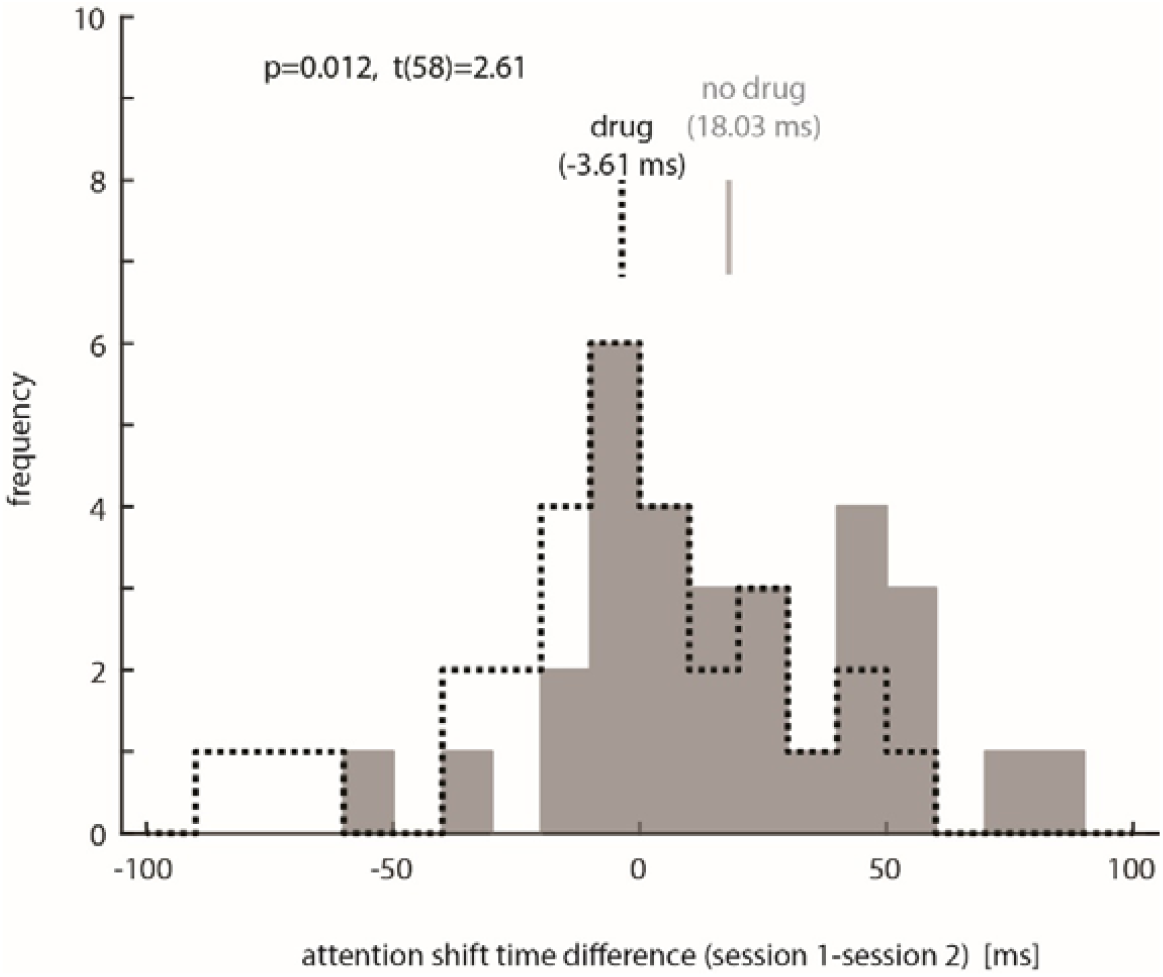
Attentional shift times differences between sessions (shift times day 1 minus shift times day 2). Grey histogram shows the distribution for the control group. Dashed black line histogram shows the distribution for the drug treatment group. Vertical black and grey bars indicate group means. Positive values indicate that shift times were longer in the first session, negative values indicate that shift times were longer in the second session. P-value (along with t-statistic) at the top indicates that distributions were significantly different.

The control group mostly had positive shift time differences, with a mean difference of 18.03 ms (Figure 6). This indicates that shift times were affected by learning in this group (becoming faster). Conversely, the group who took scopolamine after the first session showed many negative shift time differences with a mean difference of (−3.61 ms), suggesting that in this group no learning took place and shift times did not become faster. The shift time differences between groups were significantly different (t(58), p=0.012, t-test). Critically, the distribution of control subjects was significantly different from zero (p=0.0039, t(29)=3.14), while the distribution of subjects exposed to scopolamine was not (p=0.551, t(29)=-0.603). Altogether this demonstrates that shift times were subject to learning (subjects get faster between sessions), but this was blocked when subjects were exposed to muscarinic blockade during the immediate consolidation period.

## Discussion

In line with previous reports (Carlson *et al*. 2006; Chakravarthi and VanRullen 2011) we found that attention shift times were fastest for the pre-cue conditions, followed by exogenous shift, and with endogenous shift being slowest. Muscarinic receptor blockade through oral scopolamine application affected how quickly subjects can shift their attention to behaviourally relevant targets. Scopolamine increased attentional shift times in a dose dependent manner. Muscarinic challenge did not differentially affect the three cuing conditions, suggesting that the effects are similar on bottom up and top-down attention, as well as on pre-cuing conditions, which require working memory to be effective. Additionally, muscarinic receptor blockade through oral scopolamine application immediately after session 1 reduced learning induced improvements of attention shift times. Thus, muscarinic receptors are also involved in consolidation.

The contribution of cholinergic mechanisms to attention has been shown in many different settings (Nobili and Sannita 1997; Robbins 1997; Baxter and Chiba 1999; Davidson *et al*. 1999; Sarter et al. 1999; Bentley et al. 2003; Dalley et al. 2004; Herrero *et al*. 2008; Parikh and Sarter 2008; Thienel et al. 2009; Deco and Thiele 2011; Gratton et al. 2017; Dasilva *et al*. 2019), but has also often been disputed (e.g.Hangya et al. 2015). Attentional selection is assumed to be driven by frontal and parietal cortex, whereby the dorsal attention network is involved in top-down attention and the ventral network is involved in bottom-up selection (Corbetta and Shulman 2002). Parietal and frontal regions influence sensory processing directly via feedback (Moore and Armstrong 2003). How would the muscarinic signalling then be involved? Frontal areas connect to cholinergic neurons in the basal forebrain which in turn have ascending projections to sensory areas (Russchen *et al*. 1985; Sarter *et al*. 2005). If these ascending projections from the basal forebrain were relevant for attention induced improvements of sensory processing, then muscarinic blockade would dampen this (Herrero *et al*. 2008; Herrero et al. 2017; Thiele and Bellgrove 2018). But frontal and parietal areas themselves receive cholinergic input, which is relevant for attentional performance (Sarter and Bruno 1997; Parikh et al. 2008; Parikh and Sarter 2008; Gritton et al. 2016; Sarter et al. 2016; Dasilva *et al*. 2019). Reduced muscarinic receptor availability could alter overall drive in those networks, hinder competitive interactions and thereby affect how task specific populations can assemble, or how they function (Thiele and Bellgrove 2018). While our study cannot disentangle the mechanisms by which muscarinic receptors contribute to attentional shifting, it shows that different forms of attention, including working memory (pre-cue conditions) depend on adequate muscarinic drive.

Our results of muscarinic receptor contribution to consolidation and learning aligns with studies showing a cholinergic role in neuronal plasticity (McKenna *et al*. 1989; Metherate and Weinberger 1990; Weinberger and Bakin 1998; Barros *et al*. 2002) and learning (McGurk *et al*. 1988, 1991; Carli *et al*. 1997; Izquierdo *et al*. 1998; Thiel *et al*. 2002; Hasselmo 2006; Barker and Warburton 2009; Thiele 2013). Studies investigating muscarinic involvement in neural plasticity often targeted effects in sensory areas (McKenna *et al*. 1989; Brocher et al. 1992; Hasselmo and Barkai 1995; Kilgard and Merzenich 1998; Ego-Stengel et al. 2001; Weinberger 2004; Miasnikov et al. 2008). The effects seen under these conditions were not dissimilar to what occurs during perceptual learning (Karni and Sagi 1991; Schoups et al. 1995; Gold et al. 1999; Schoups et al. 2001; Pleger et al. 2003; Li et al. 2005; Sanayei et al. 2018), an implicit type of memory. Other types of implicit learning, such as conditioned avoidance, cue triggered reward, and motor learning, depend at least partly on cholinergic signalling and plasticity of the basal ganglia (Packard and Knowlton 2002; Ostlund et al. 2017). Whether the learning seen in our study is related to perceptual, cognitive, or covert motor learning is unclear. Whatever the source, its consolidation is dependent on muscarine receptor availability.

## Conclusions

We found that muscarinic blockade affected attention shift times in a dose dependent manner. No differences were found between effects on endogenous, exogenous, or pre-cued attention shifts. Moreover, application of the muscarinic blocker scopolamine during the immediate consolidation period resulted in reduced improvements of attention shift times.

## Acknowledgments

AT was funded by Wellcome Trust 093104 (AT), and the MRC MR/P013031/1.

